# Invariant Object Representation Based on Principle of Maximal Dependence Capturing

**DOI:** 10.1101/662130

**Authors:** Rishabh Raj, Dar Dahlen, Kyle Duyck, C. Ron Yu

## Abstract

Sensory inputs conveying information about the environment are often noisy and incomplete, yet the brain can achieve remarkable consistency in object recognition. This cognitive robustness is thought to be enabled by transforming the varying input patterns into invariant representations of objects, but how this transformation occurs computationally remains unclear. Here we propose that sensory coding should follow a principle of maximal dependence capturing to encode associations among structural components that can uniquely identify objects. We show that a computational framework incorporating dimension expansion and a specific form of sparse coding can capture structures that contain maximum information about specific objects, allow redundancy coding, and enable consistent representation of object identities. Using symbol and face recognition, we demonstrate that a two-layer system can generate representations that remain invariant under conditions of occlusion, corruption, or high noise.

## Introduction

The world is organized into objects. The goal of sensory processing is to generate representations of the objects in cognitive centers that allow detection of meaningful association among them (1). For individual objects, a suitable representation must permit precise identification and facilitate cognitive robustness by remaining consistent for its various input forms (2). While individual features shared among multiple objects cannot be used to unambiguously identify any object, structural relationships among multiple features can provide information to uniquely characterize them in diverse situations (3–5). For example, a predator can be recognized in various concealments or camouflages because the relative configuration of its features, though only visible partially due to concealment, provides sufficient information for it to be identified. Any sensory processing framework, therefore, must address how structural relationships among features that are most informative about objects can be encoded along the sensory pathways to generate object representations that remain consistent across different experiences.

Perceptual invariance requires representations to be stable for the same objects yet sensitive enough to distinguish different ones (6, 7). Finding a basis of representation that satisfies these two diametrically opposed requirements remains a fundamental challenge to any computational model of sensory processing. Most current models are based on two theoretical foundations-efficient coding (8, 9) and hierarchical assembly (9–12) -- to build representations. Adopted from Information Theory, efficient coding has focused on minimizing redundancy in information transmission by encoding independent features present in natural stimuli (13–17) (**Fig. 1A**). However, this principle appears only applicable to early processing stages. Complex tuning characteristics found in high cortical areas are explained by hierarchical assembly, which progressively combines the independent components to generate complex features to represent the objects (18–21). Various models have been proposed to account for perceived invariance in location, scale, orientation, and rotational views of objects (22–29). Whereas these models can account for novel viewpoints or forms (21, 25–31), they are not effective in mapping corrupted or occluded forms of objects to the same representation (**Fig. 1B**). Although independent features with minimal redundancy allow efficient information transmission (16, 17, 32, 33), the process of combining them into more complex forms cannot effectively compensate for lost features from the input. To compensate for information loss, mechanisms of inference (e.g., pattern completion) must be evoked (30, 31). This not only introduces a new set of computational rules separate from that involved in processing primary sensory input, but also violates the redundancy reduction principle.

**Figure 1.**
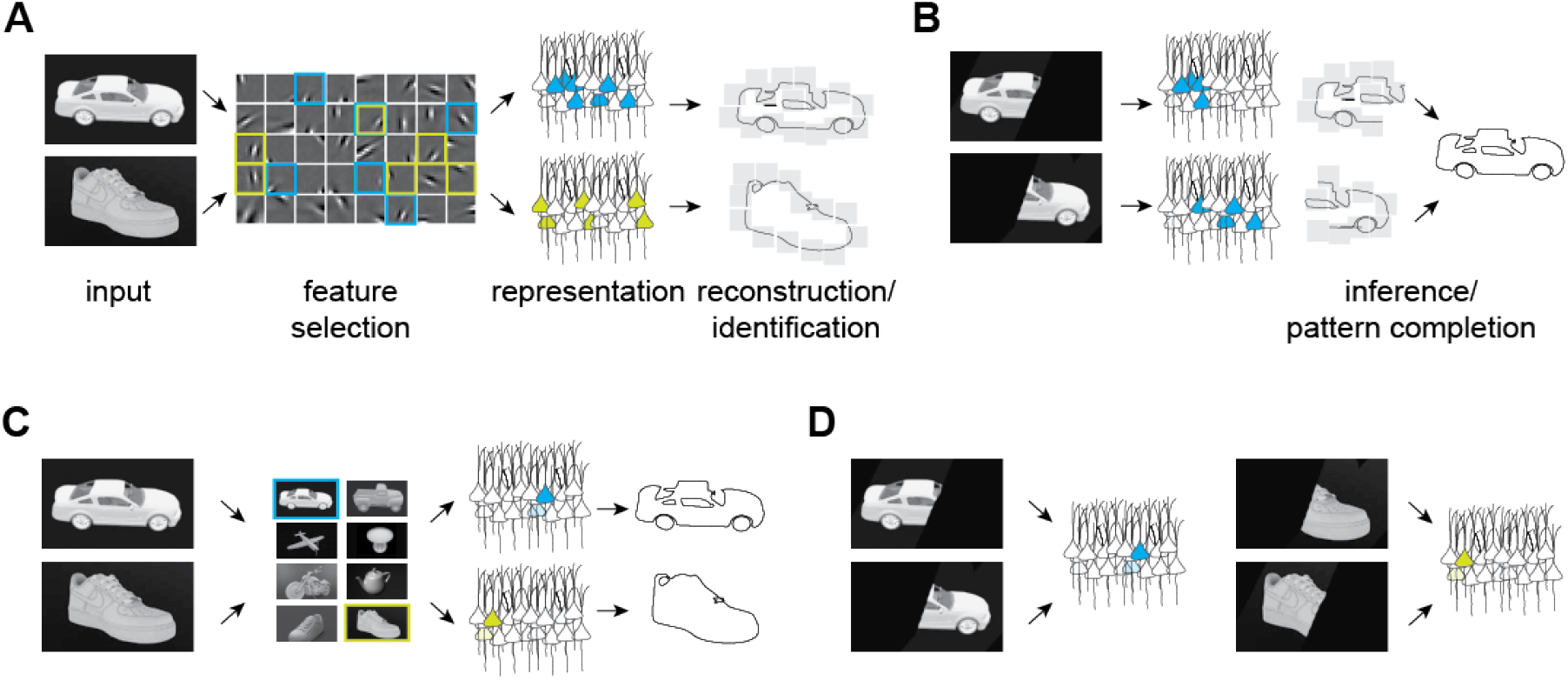
Illustration of the classic and the maximal dependence capturing frameworks for object representation. **(A),** Classic framework. Inputs are decomposed into independent features and represented by neurons (color-coded for separate objects). Object identities are determined by the composition. Gray patches depict the receptive field properties of representation neurons. **(B),** Occlusion, or missing features presents a problem for the classic framework because they cannot be recovered through transformation of input signal into the same representation. Additional computation is needed for identification. **(C),** In the MDC framework, coding units capture structural relationships among the features and encode them as a whole. **(D)**, Since the features are redundantly encoded, missing feature does not affect encoding of the identity of the objects.

Recent studies have attempted to address perceptual robustness by relating models of deep learning, such as convolutional and recurrent neural networks, to sensory processing (34, 35). Although it has been suggested that these network models can recapitulate aspects of sensory processing, all models require extensive training using a large number of examples to achieve robustness, and often require artificially high numbers of network layers. In contrast, the biological brains, regardless of their size and sophistication, can recognize objects in the presence of noise or corruption without deep layers or learning from a large number of corrupted examples. It is not clear whether the deep learning frameworks recapitulate the computations employed by the brain (36).

The identity of an object is established by the structural relationship among its features. As associations among features uniquely define an object, capturing these associations could lead to robust coding of object identities. Thus, we propose that sensory processing should follow an alternative principle from redundancy reduction and aim to capture as many as possible associations among features that are informative about object identities to maximize mutual information between the coding units and individual objects. We refer to this as the maximal dependence capture (MDC) principle. The principle allows repeated coding of the same features in different configurations and preserves information about their structural relationships. Missing elemental features can be inferred from their dependence with others, thus allowing redundancy to enhance robustness against signal corruption. In his more recent work, Barlow has suggested redundancy is useful for encoding object identities, but it is not known how to implement it (37). In this study, we establish a mathematical framework and demonstrate its robustness in transforming corrupted or occluded images into consistent representations without the need of deep layers, large training sets, or training on corrupted examples. Importantly, this framework is adaptive according to the object sets. It can lead to simple receptive field properties as found in early processing stages, as well as complex receptive fields as those found in higher order centers. These findings suggest that a single set of computational rules can account for both primary and high order processing of sensory information.

## Results

All features are not equally informative about the identity of an object. Features or feature combinations unique for the objects are the most informative. A coding strategy relying on independent features does not serve to capture the informative elements. To be effective, a coding strategy should encode highly informative structures corresponding to the identity of an object (**Fig. 1C**). At the extreme, a single coding unit encodes the intended object as a whole by capturing all features that define the object. This creates a scenario where each unit encodes a distinct object such that partial inputs that carry sufficient information can overwhelmingly activate it to make the representations invariant against input corruption (**Fig. 1D**). Whereas this simplistic paradigm has been used to encode highly specific stimuli, it is not known whether it broadly applies as a general principle of object representation. To be effective, such coding paradigms should have the ability to capture most structural relationships while encoding large numbers of objects.

We have derived a coding strategy based on a dimensionally expanded pattern transformation that enforces sparsity in representing objects. Although similar transformations, particularly the sparse coding transformation of Olshausen and Field (17, 33), have been used to learn independent features from natural environment, they tend to generate arbitrarily complex receptive fields that do not naturally occur in the input (38). To overcome this problem, a non-negative constraint is applied in the learning process. Another important distinction of this transformation is that it is not intended to identify independent components in the stimuli based on the statistics of the entire input space. Rather, it is adaptive, i.e., it is optimized for a limited set of inputs to capture the most informative elements. These characteristics, as we show below, are important to generate robust representations.

For concreteness, we consider a system with *M* input and *K* output units to encode *N* objects. Each object is manifest as a specific pattern of the input and output units. The encoding problem is to find a basis set or a dictionary (***Φ ∈ R***^M x ***K***^) that relates the matrix of input signals (***X ∈ R***^M x ***N***^) to a matrix of object representations (***A ∈ R***^K x ***N***^). We treat the learning process as the matrix factorization ***X*** = ***ΦA*** with an objective function that minimizes the *Frobenius* norm to preserve information and the *l_0_* norms (implemented as minimizing *l_1_* norm) to maintain sparseness, i.e.,

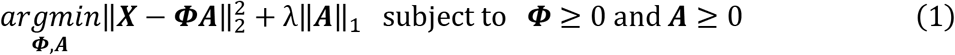

Here, *K* is set to accommodate as large an *N* as possible to maintain sparseness. It is assumed that *K* will exceed *M* in most cases such that the transformation is dimensionally expanded. Non-negative constraint is applied to every iteration during the optimization steps (see methods). With a learned dictionary, an optimal representation for any input pattern is obtained by solving the optimization problem:

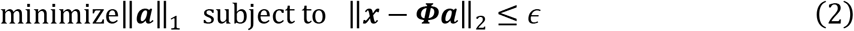

The problem utilizes sparseness to uniquely solve an under-determined system of equations. Taken together, learning a dimensionally expanded dictionary from a finite set of objects with a non-negativity constraint, distinguish this transformation from other approaches that use direct convolution of input with the dictionary to generate representations (30, 33, 39).

### Dependence capturing

We illustrate key characteristics of this framework by encoding of binary symbol images from world languages, which also allows us to quantitively measure the robustness of the coding framework (**Figs. 2, S1A**). With 256 input and 800 output units, this is a dimensionally expanded representation (**Fig. 2A**). Learning to represents 1000 symbols results in dictionary elements that contain both local and global structures (**Fig. 2B**). The representations of symbols are sparse (**Fig. 3**). The structures of the dictionary elements, which can be considered as equivalent to the receptive field properties of neurons, resemble the symbols themselves, thus meeting the goal of capturing structural relationships that uniquely characterize inputs. Because the symbol set contains images that are similar, multiple elements share similarities to each other. For example, several dictionary elements shown in Fig 2B share oval-like shapes. The high similarity among dictionary elements indicates that the same set of elemental features are redundantly encoded as required to capture most of the input structures (**Fig. 2C**). To quantify the similarity between dictionary elements and the objects, we calculated the activation probability of individual pixels from the input set and from the set of dictionary elements. The two are closely matched (**Fig. 2D**) and have nearly identical bit entropy (**Fig. 2E**). These results illustrate that dependence among the input units is maximally captured by the dictionary elements of the encoding layer. The computation in the MDC framework forces the dictionary elements to extract structural relationships among pixels that are naturally present in the stimuli without creating arbitrary or overly complicated dictionary elements.

**Figure 2.**
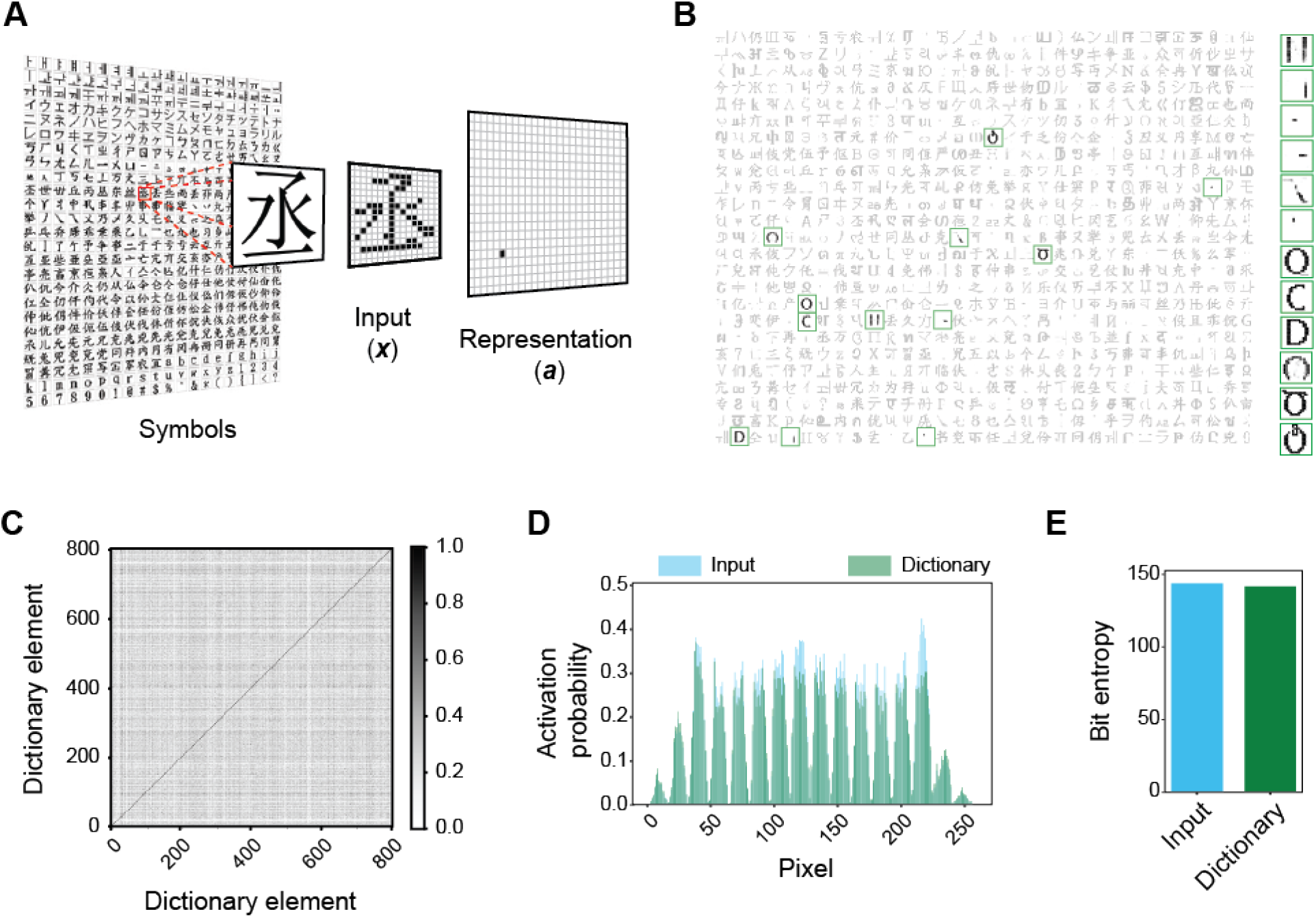
Dependence Capturing by the MDC framework. **(A),** Illustration of encoding in the MDC framework. Symbols from world languages are converted to 256 (16×16) pixel images (***x***) that are transformed into the activity of a set of 800 representational units (***a***) to encode symbol identities. An example of the process is shown using a character encoded by a single representational unit to indicate the ability to encode complex structural features. **(B)**, Structures of the dictionary elements (receptive fields) learned from 1000 symbols. Highlighted elements that are displayed in larger size are the ones most active by the inputs shown Figure 3. Note the similarity among some of the elements. **(C)**, Similarity between individual dictionary elements, calculated as cosine similarity, is high. **(D)**, Probability of activity of individual pixels in the input vs. dictionary space. The two exhibits high similarity. The periodicity reflects that the pixels at the edge of the frame do not have activations. **(E)**, Bit entropy of input and dictionary elements.

**Figure 3.**
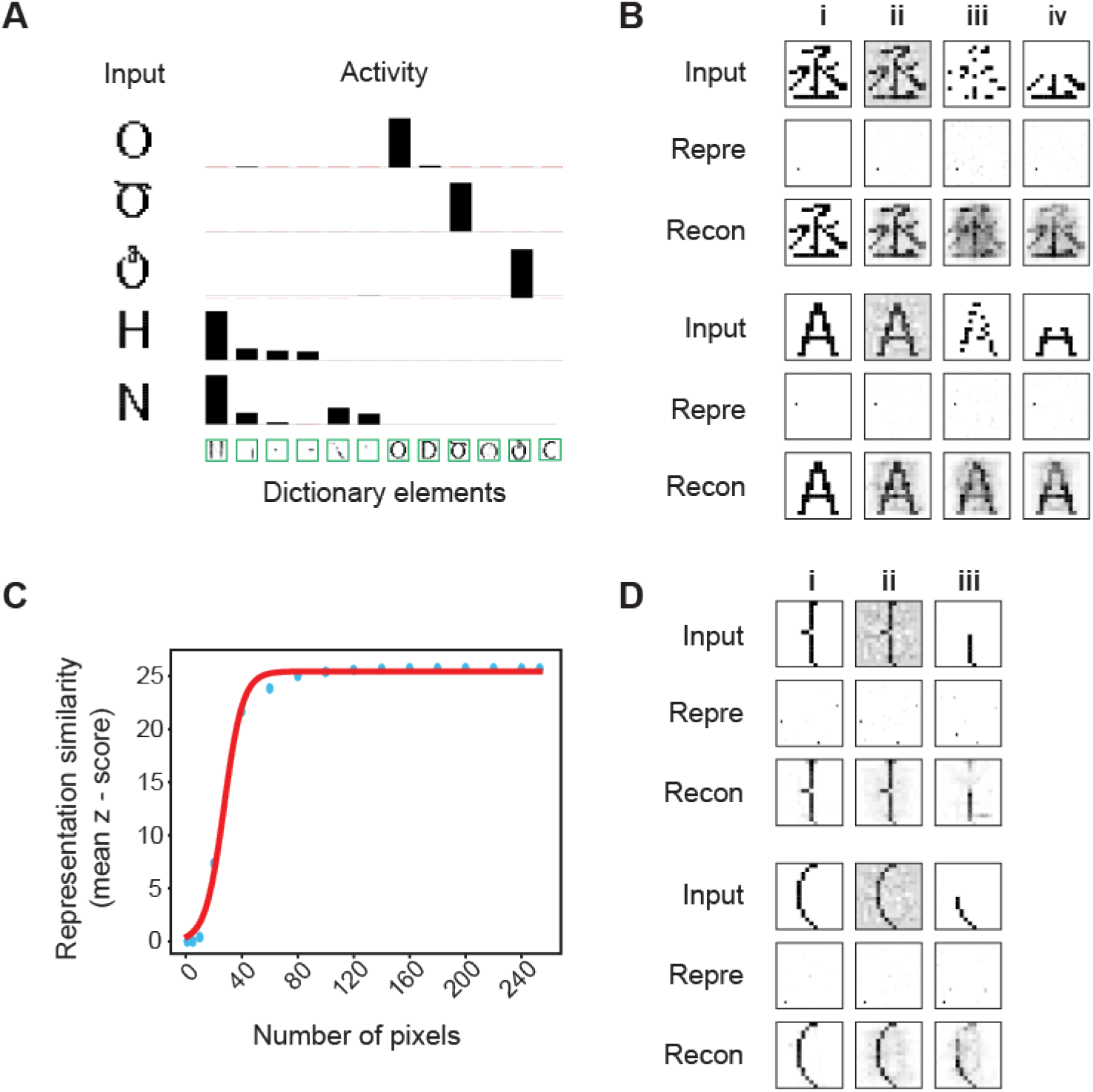
Invariant object representation. **(A),** Representation of inputs by activity of the dictionary elements. Height of the bars indicates activity evoked by different input patterns. Only the most active ones are show. Note that highly similar inputs only activate the dictionary elements that are most similar to them despite the similarity among the elements. **(B),** Examples of coding of the original (**i**) and corrupted input under noisy **(ii)**, pixel loss **(iii),** and occlusion **(iv**) conditions. Representation of the corrupted signals (Repre) are similar to those of the originals. Reconstructed images (Recon) from the dictionary elements resemble the original symbols. **(C),** Z-scored similarity between representations of corrupted and original signals as a function of total pixels in the input layer (randomly selected as in **B iii**). Scores calculated with different pixel numbers (blue dots) is fit to a sigmoidal curve (red line). **(D),** Example of two highly similar symbols being distinctly and robustly represented. The original input signals (**i**) corrupted by noise (**ii**) or occlusion (**iii**) are transformed to output activities that are similar to each other. Reconstructed images recover the original signals.

### Invariant representation of corrupt input signals

We next test whether the MDC framework can encode objects distinctively, especially in conditions of noise and corruption. In classic frameworks, neural network models require deep layers to enhance robustness. The MDC framework captures high order structural relationships from the inputs, thereby can be robust against corruption without deep layers. The two-layer model based on our framework readily distinguishes highly similar patterns and represents them differently (**Fig. 3A**). Representations of the symbols are sparse (**Fig. 3B**), and there are similar non-zero correlations among pixels in both the input and dictionary (**Fig. S1B**). The correlation matrix of output units is very close to identity (**Fig. S1C**).

Moreover, this framework presents a solution to the invariant representation problem. Without learning from corrupt examples, the model can transform input signals corrupted by Gaussian noise (**Fig. 3B, ii**), randomly missing pixels (**Fig. 3B, iii**), or partial occlusion (**Fig. 3B, iv**) into representations nearly identically to those of uncorrupted signals (**Fig. 3B, i**). Reconstructions of the images using dictionary elements resemble the whole symbols rather than parts (**Fig. 3B**). Using Z-score of the pairwise cosine distances between the representation of corrupted input and all learned symbols, we observe high specificity for the correct input-symbol pairs, indicating that the framework generates highly specific representations. In Monte Carlo simulations with randomly missing input units, representations with high specificity are obtained with as few as 60 (23.4% of the 256) units (**Fig. 3C**). Importantly, the decoding algorithm is sensitive to small differences in the input patterns. For example, two highly similar input patterns are represented differently and robustly under various corruptive conditions (**Fig. 3D**). Thus, the computational framework fulfills the requirements of stability and sensitivity set forth by Marr and Nishihara (6) without requiring deep layers or learning from a large number of variable examples.

### A robust code for face recognition

We next tested the two-layers model in encoding complex, non-binary signals such as human faces (**Fig. 4**). We trained the model on 2000 human faces (**Fig. 4A**). The learned dictionary elements are neither parts of faces nor do they resemble whole faces. Instead, they are composed of complex assemblage of facial features, again suggesting that the algorithm captures structural relationships among the features present in the training set (**Fig. 4B**). Representations of the faces are stable, unique and robust against common alterations such as headwear, facial hair, or eyewear **(Fig. 4C)**. The same person is represented nearly identically when a mustache, a pair of sunglasses, or both were added. Even when half of a face was blocked in different positions, the framework produces the same representation (**Fig. 4D**). Inversely reconstructed images from the representations are similar to those of unadulterated faces even when the faces are half blocked (**Fig. 4C and D**). Notably, the robustness is achieved from learning only 2000 examples, and without using any corrupted images. This is in direct contrast to many approaches using large numbers of variegated examples as training sets. Moreover, the dictionary learned from the training set can be applied to a completely new set of faces. We used it to obtain representations of facial images in the Yale face base, which contains 15 individual faces 11 different lighting conditions and facial expressions. The representation correctly categorized the faces according to the individuals (**Fig. S2**)

**Figure 4.**
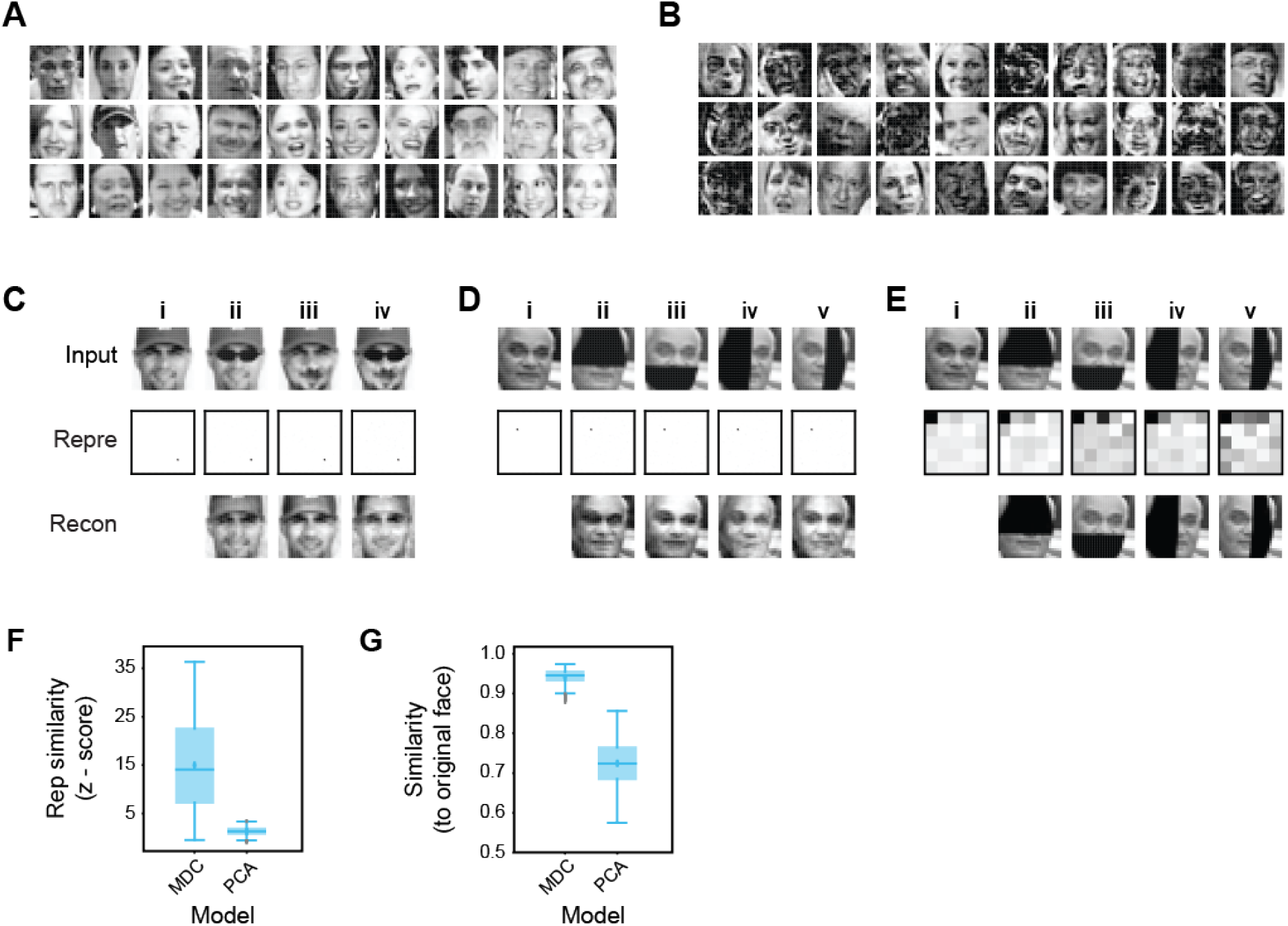
Robust representation of human faces. **(A),** Examples of face images in a library of 2000 (1000 male and 1000 female) faces used to train a two-layer model. **(B),** Examples of dictionary elements learned from the face library. Note that they incorporate complex combinations of facial features but are not necessarily part or whole of any specific face. **(C),** Representation and recovery of faces with different alterations. A face (**i**) was altered to wear a mustache (**ii**), a pair of sunglasses (**iii**) or both (**iv**). Representations of the altered faces were nearly identical to the original even though these examples were not in the training set. Images reconstructed from the representations based on the dictionary were similar to the original images. **(D),** Representation and recovery of occluded faces. Four different occlusions of a face - top (**ii**), bottom (**iii**), left (**iv**), and right (**v**) were generated. Representations of the occluded faces were highly similar to the original one. Reconstructions also matched the original face. **(E),** Face identity was not preserved in representations based on PCA performed on the 2000 training faces. Representations of original (**i**) and occluded faces (**ii-v**) were obtained in the principal space (the first 25 components are shown to highlight the differences). Representations of occluded faces are different from original. Reconstructed images match the corrupted rather than original faces. **(F-G),** Specificity (Z-score) of face representations **(F)** and similarity (cosine distance) between reconstructed and original images **(G)** were calculated for MDC and PCA models. 50 faces were chosen randomly from the training set to create the four occluded versions.

We compared our code against a recently proposed, principal components based face code (40). Using dictionaries obtained through principal component analysis (PCA), the same face with different parts occluded generated different representations. Image recovery produces occluded, but not uncorrupted images (**Fig. 4E**). Quantification of specificity using Z-scores shows that our model generates representations that are highly specific in matching the original face (**Fig. 4F)**.

Recovered images from representations of corrupted inputs are highly similar to the originals (**Fig. 4G**). PCA-based decoding does not exhibit such selectivity or similarity (**Fig. 4F and G**). Thus, combinatorial face code based on the MDC framework is robust against corruption whereas the one proposed before is not (40, 41).

### Optimal decoding using l1 minimization

By capturing dependencies in the input signals, the dictionary elements can have large overlaps. The output units corresponding to these dictionary elements are expected to have high correlations in their activities in response to the inputs. This expectation appears counter to the prevailing notion that the encoders should be decorrelated (10, 32, 42, 43). The contradicting expectations are reconciled by the *l_1_* minimization process. Convolving the input with the dictionary results in initial broad activation of the encoding units (**Fig. 5A**). However, once *l_1_* minimization is applied, activity is restricted to a few units (**Fig. 5A**). Thus, *l_1_* minimization facilitates the decorrelation of activities. Indeed, the stability of our computational framework in generating invariant representations relies on *l_1_* minimization, which can find the near-optimal solutions (i.e., closest to *l_0_* solutions) for sparse signals (44–46). Classic linear models such as non-negative factorization (47) or non-linear input transformations are not sufficient to generate representational stability. We compared *l_1_* minimization against Locally Competitive Algorithm (LCA) and the classic Linear Non-Linear (LNL) approaches in generating representations of occluded faces. *l_1_* minimization produces representations with the highest specificity (**Fig. 5B**). Although LCA performs better than the LNL, neither produces representations with the specificity of the *l_1_* minimization algorithm. LNL does not produce sparse solutions, whereas LCA produces a solution that only utilizes a subset of components derived from *l_1_* minimization (**Fig. 5C**).

**Figure 5.**
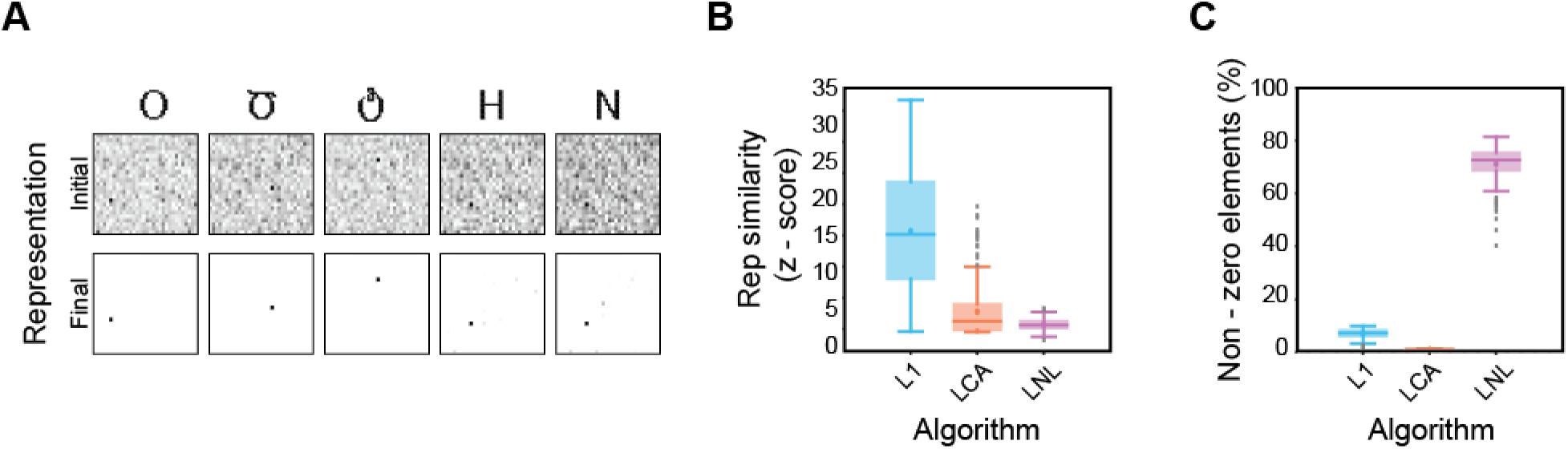
Near-optimal sparse solutions are necessary for high specificity decoding. **(A),** *l_1_* minimization ensured sparsity permits distinction among similar objects. Representation of five symbols. The initial activity is driven by convolving input with dictionary elements before *l_1_* minimization. Final activity is after *l_1_* minimization. **(B),** Comparison of specificity among three algorithms in obtaining sparse representations of 50 occluded facial images as in Fig. 4D and 4E. *l_1_* minimization method produced representations with the highest specificity. **(C),** Comparison of sparsity of the representations (measured as the number of non-zero components in the representations) obtain through the different methods. Representations from *l_1_* minimization are optimal and sparse. LCA recovers more sparse representations because it only recovers specific components of the optimal solution. LNL does not produce sparse representations.

### Adaptive nature of the MDC code

In the examples above, individual objects are represented by very few active units and the dictionary elements are complex. Whereas complex receptive fields are observed in high order brain areas, particularly those involved in encoding objects and faces, it is uncommon to have extremely sparse responses. Can the MDC model recapitulate experimental observations? In biological brains, the number of neurons is constrained by evolution and anatomical structures and cannot increase indefinitely. Although we have set *K* to be large to accommodate large numbers of objects, an increase of the number of objects *N* is expected to influence the code structures. We explored this relationship by setting *N* and *K* at different values. With *N* fixed, at low *K*, dictionary elements are more localized (**Fig. 6A**). The response becomes less sparse, and it requires more active encoders to encode the same symbol (**Fig. 6B**). An increase in the number of symbols to be encoded causes a divergence of the dictionary element structures from the input patterns, which can be measured using the K-L divergence (48) of pixel distribution between the input and the dictionary (**Fig. 6C**). Redundancy for the output units also increases with increasing number of symbols (**Fig. 6D**). At high *N* numbers, the redundancy approaches that of the input, suggesting the code has become less efficient. The same observation is made for complex signals. At low *K* values, the dictionary elements for faces are more localized, resembling local features of faces (**Fig. 6E**). At high dimensions, they become more complex and face-like (**Fig. 6F**). More units are required to encode each face at low dimension (**Fig. 6G**).

**Figure 6.**
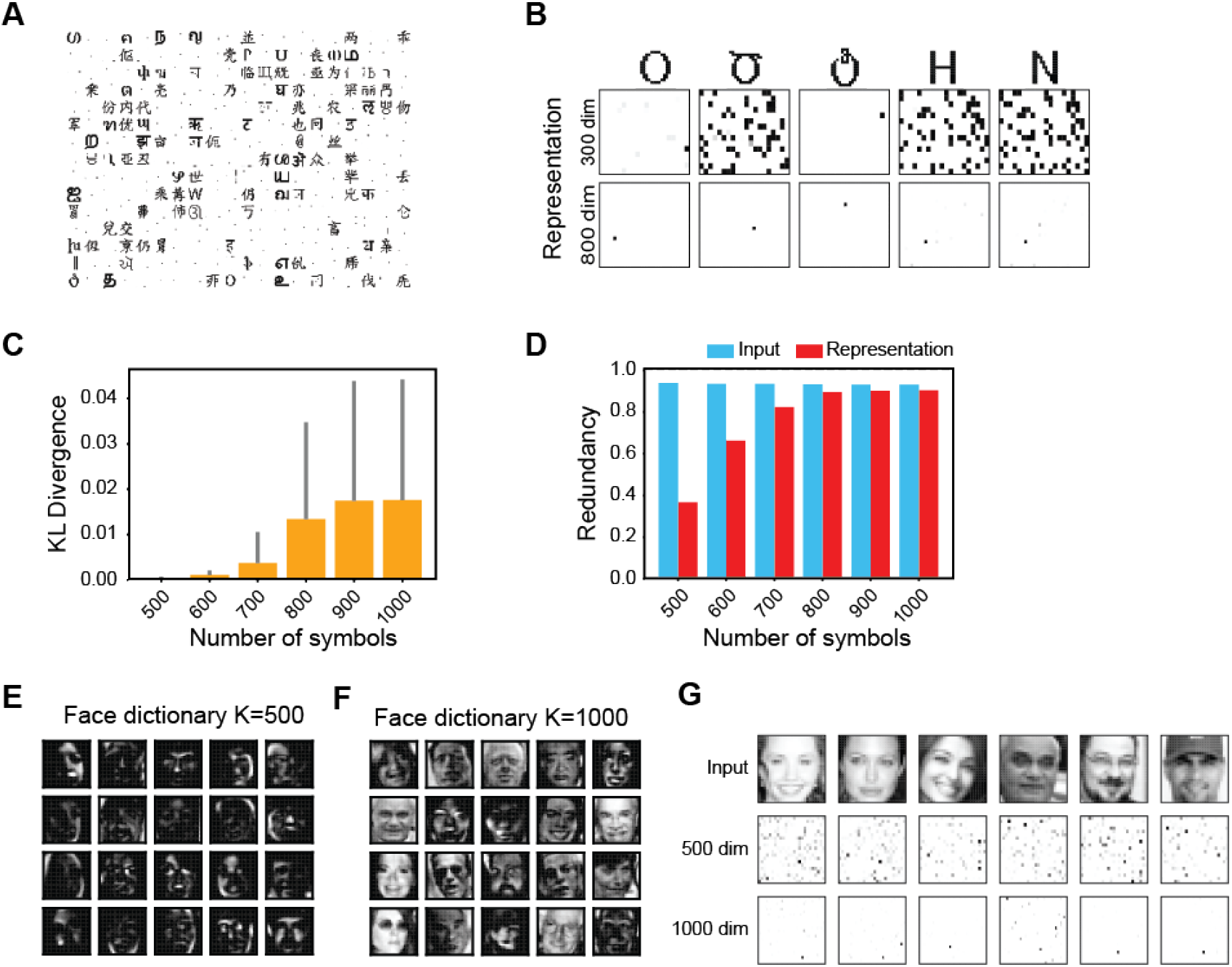
Sparsity of responses determined by the number of objects and dimension of the encoders. **(A),** The structures of dictionary elements for the symbols in Fig. 2 with a 300-unit output layer. Compared with that of the 800-unit shown in Fig. 2B, there are more localized features. **(B),** Representation sparsity increases with increased dimensions. **(C),** K-L divergence between pixel distributions in the input signal and in the dictionary element as a function of the number of symbols to be encoded. **(D),** Coding redundancies in input and output (representation) while encoding increasing numbers of symbols. **(E and F),** dictionary elements of faces with a 500-dimension (**E**) or 1000-dimension (**F**) encoder-set. **(G),** Response elicited by faces are sparser with increased dimensions.

The premise of the MDC framework is that individual encoders maximally capture structural dependencies in the input signal, which results in complex dictionary elements. This appears to be contradictory to the localized receptive fields observed in the primary sensory cortices, which have been explained by models that capture independent components of natural inputs based on the Efficient Coding hypothesis. Is the MDC framework able to produce these localized receptive fields?

That more localized dictionary elements are observed with large *N* suggests that a large number of stimuli may force the neurons to tune to more localized features. In the visual system, all visual information is relayed through the primary visual cortex, i.e., the primary visual cortex must accommodate all visual objects. The overwhelming number of visual objects could force the cortical cells to tune to more localized features in visual signals. To test this hypothesis, we trained the network with varying numbers of image patches taken from natural images as individual stimuli (49) (**Fig. 7A**). This is similar to the approach of Olshausen and Field (17), but with non-negative constraint. To accommodate this constraint and simulate the On and Off channels in the mammalian visual system, we parsed the images into positive and negative signals. With training using up to 30,000 patches, both local, simple cell like dictionary elements as well complex elements have developed (**Fig. 7B**). With low number of training images, the dictionary elements are relatively complex (**Fig 7C**). Despite high correlations among the images, encoding units are highly decorrelated (**Fig. S3A and B**). Increasing the size of the training set gradually produces dictionary elements with localized and orientation-selective projective fields that are similar to the receptive fields simple cells in the mammalian primary visual cortex (19) (**Fig. 7B and C**). The simple cell like dictionary elements resemble Gabor filters as found in earlier studies (32, 50). Interestingly, complex projective fields also persist in all training conditions. The percentage of simple projective fields in the dictionary increases with the size of the training set. With a fixed set of training images, an increase in the encoding dimension reduces correlation among the encoding units but increases the correlations among dictionary elements (**Fig S3C**). Thus, similar to classic interpretation, simple receptive fields emerge when large numbers of natural images are encoded under the MDC framework (13–15, 43, 51). Importantly, complex tuning is always present in our model without requiring it to be synthesized from simple cells as the classic model predicts (19).

**Figure 7.**
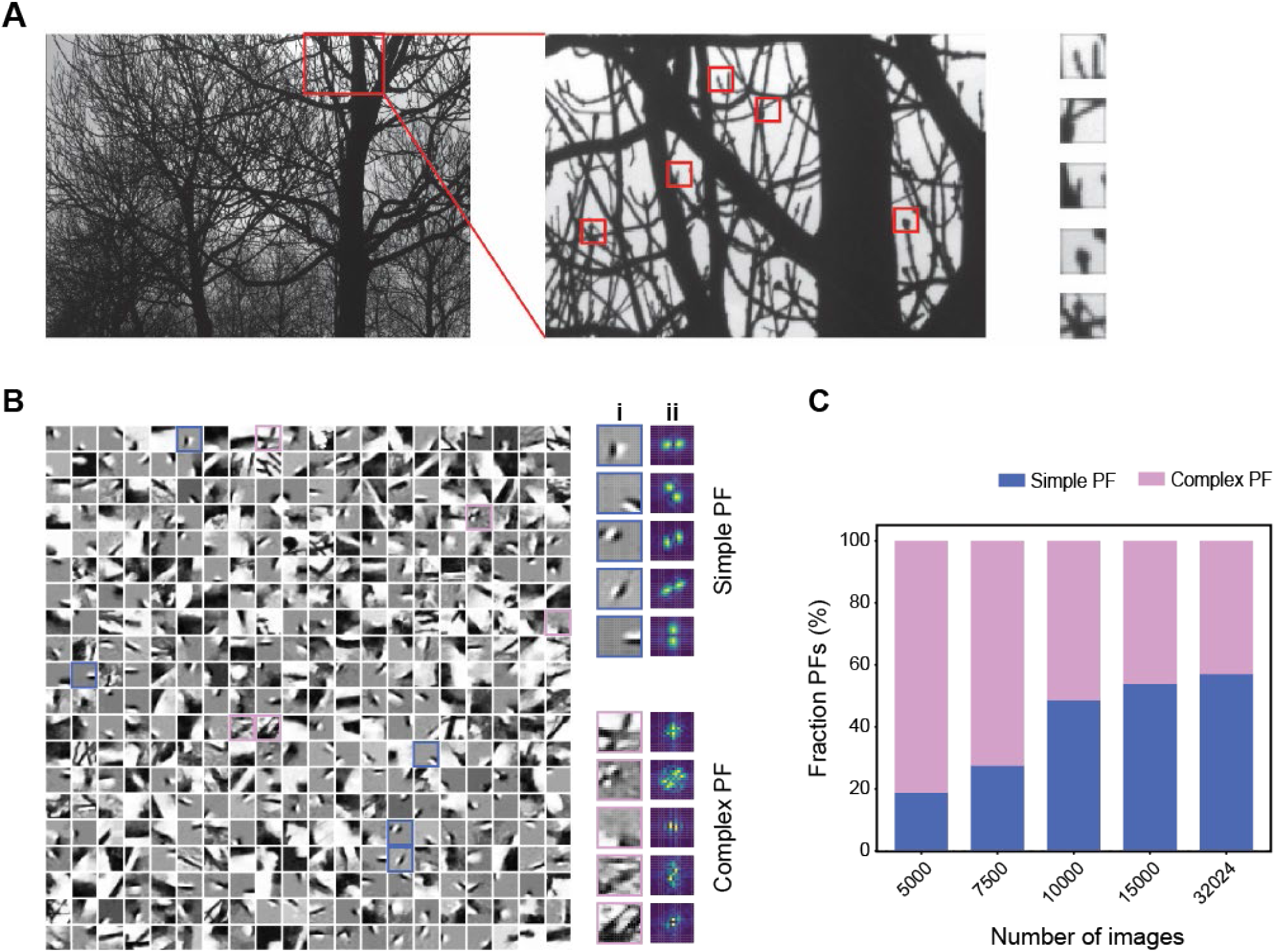
Emergence of simple and complex receptive fields from training with natural images in MDC. **(A),** Illustration of image patches derived from natural scene. **(B),** Examples of structures of individual dictionary elements. Both simple (cyan) and complex (magenta) elements are observed (**i**) and can be determined using Fourier transforms of the structures (**ii**). **(C),** As the number of training images increases, the tuning properties of encoders become more localized and the percentage of simple PFs increases.

## Discussion

We propose that maximal dependence capture is a general principle in sensory processing as an alternative to the redundancy reduction principle. The prime motivation behind redundancy reduction is to arrive at a factorial code for object representation (1, 9). However, a factorial code based on independent features is not suitable for invariant coding, especially in cases involving corruption or occlusion. Hierarchical models based on feature combinations can be utilized to learn feature associations, but it requires learning of multiple instances of feature combinations originating from the same object in order to achieve robustness. It also disrupts the factorial nature of representations. The MDC framework resolves this complication by letting the coding units to extract the most informative combination of features that uniquely identify specific objects. Capturing maximum dependence among features in a sparse code preserves the combinatorial nature of the code while enabling the system to utilize the dependence to achieve robustness in object identification.

The sparseness in representations suggests that the code is efficient in encoding object identities. However, as feature combinations that uniquely define objects are not necessarily independent, efficiency in encoding stimulus features is less in the MDC framework than what could be achieved by the classical framework. Arguably, from the evolutionary perspective, the sensory systems have evolved to detect ethologically relevant signals. Analyzing environmental stimuli and parsing them into components of minimal redundancy is not necessary for this goal. In this regard, capturing unique associations among features enables the system to rely on redundancy to achieve robustness in object identification with a minimal compromise in coding efficiency.

Sparse representation of objects is also reminiscent of the “grandmother cells”. These cells have been considered as computationally implausible (52), yet neurons have been recorded to respond to individuals regardless of the form being presented (53, 54). The improbability of constructing a grandmother cell, we argue, is based on the hierarchical assembling of an invariant representation from independent components, which can create the problem of combinatorial explosion to account for all possible combinations of features to generate singular representations. In the MDC framework, no such hierarchical assembly is required. In fact, the generation of “grandmother cell” like representation is a feature of the framework.

We have proposed a mathematical model that conforms to the MDC principle. It achieves dependence capturing by seeking maximal sparsity in a dimensional expanded representation. Dimension expansion is found in many sensory systems. In this model, the combination of non-negativity and sparsity constraints allows the effective capture of naturally occurring associations among morphological features while rejecting spurious ones (47). Sparsity forces the encoding units to tune to distinguishing groups of morphological features of the stimuli such that the fewest units can capture the information about the source. The non-negative constraint, while serving as a proxy of neuronal responses carried by spikes, prevents the use of negative coefficients to subtract features from arbitrarily complex dictionary elements. Thus, the framework forces the dictionary elements to extract structural relationships among pixels that are naturally present in the stimuli without creating arbitrary or overly complicated dictionary elements. We note that past approaches have sought sparse codes, using overcomplete basis set, or impose non-negativity constraints (17, 33, 55, 56), but those models do not achieve the robust invariance in dealing with corruption and occlusion demonstrated here. Our results show that both non-negative constraint and minimal *l_0_*-length sparse codes are required to effectively capture the structural relationships in the objects. As we demonstrate in the study, simply obtaining a sparse code is not sufficient to render the representation robust against corruption. We do not know whether and how specific forms of computations are implemented in a neuronal network to achieve it *l_1_* minimization-like computation. It is likely implemented in the nervous system as lateral inhibition. We suspect that a combination of divisive normalization (57) with a sparseness-driving mechanism could achieve the goal.

A unique characteristic of the MDC framework is that the receptive fields of individual encoding unit is expected to capture complex structures embedded in the objects. This requirement creates a situation that the units can be highly similar in their tuning properties. This characteristic may appear antithetical to the notion of efficient coding, which demands individual encoding units to be as independent as possible. Our results indicate that these two characteristics are not contradictory. Indeed, the responses units are highly decorrelated even though their receptive fields have high levels of overlap. This is because the algorithm can resolve the competing activities and lead to sparse, decorrelated response. Moreover, physiological support for such structurally complex receptive fields of neurons can also be found in a recent study which has shown that cells in the inferotemporal cortex of monkeys respond complex combinations of structures from objects (58). In addition, as we show, both simple and complex receptive fields can be produced by the same model. This highlights the relationship between the number of distinct objects to be encoded and the number of available coding units.

In its sparsest form, the receptive fields of individual coding units resemble the objects themselves. One may consider this as a form of template matching. As we have shown, however, the MDC framework is not template-matching. The captured receptive fields may resemble whole structures of the objects, but they are not identical. Moreover, these receptive fields change as the coding units adapt to different numbers of inputs.

Maximal dependence capturing, therefore, offers a single framework to explain seemingly contradictory observations on receptive field properties of neurons in different hierarchies in the brain. It allows the encoding of the features and their relationships in the same step. Importantly, the MDC framework enables invariant representations. Because of the ability to effectively capture the structures in the signals, models built under this framework do not require learning from a large number of examples, nor from corrupted signals. It also does not require deep structures to achieve robustness. On the other hand, deep structures are likely to further improve coding efficiency and robustness.

## Acknowledgments

**General:** We thank Drs. K. Si, J. Unruh, P. Kulesa, M. Klee and members of the Yu laboratory for insightful discussions. This work fulfills, in part, requirements for RR’s and KD’s Ph.D. thesis with the Open University, United Kingdom.

## Author Contributions

C.R.Y conceived the idea and supervised the research. C.R.Y and R.R. developed key concepts and co-wrote the paper. R.R and D.D. performed analyses and modeling. K.D. generated face database.

## Materials and Methods

### Learning algorithm

Dictionary learning is treated as a blind source separation (BSS) problem (59, 60). An input signal is modeled as the response of *M* primary encoders. In the case of images, *M* = *m_1_* · *m_2_*, where *m_1_* and *m_2_* are the horizontal and vertical dimensions of the images. A set of N signals is presented for training as an input matrix 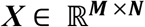, representing the response of *M* pixel to *N* patterns. The matrix ***X*** is then factorized into two matrices ***A*** and ***Φ***, so that ***X*** = ***ΦA***. Here, 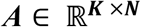 is the matrix representation of *N* patterns in a ***K*** dimensional basis set defined by 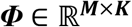.

To get the factor matrices through BBS, we imposed restriction on ***A*** to be sparse. The measure of sparsity was chosen to be *l_0_* norm, but the solution is achieved through minimizing *l_1_* norm. In addition to this sparsity constrain, we demanded both ***A*** and ***Φ*** to be non-negative.

Several possible BSS algorithms could result in an appropriate matrix decomposition under the given constraints (56, 61, 62). In particular, we used non-negative blind source separation algorithm nGMCA (61, 63, 64). When a *l_1_* measure of sparseness is used, then the sum of the absolute values of coefficients of ***A*** is minimized. The minimization problem takes the form of:

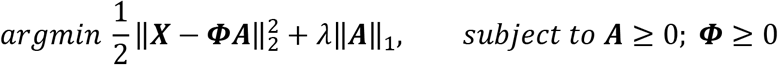

Thus, the process to solve this problem requires the minimization of the *Frobinius* norm difference (i.e., the Euclidean Distance) between the two sides of the equation and the minimization of the *l_1_* norm.

Each time BSS is performed, the ***Φ*** matrix was seeded with random numbers. Optimization was performed until convergence or when predefined number of iterations was reached.

### Sparse coding

Once the dictionary ***Φ*** is learned, any input pattern can be transformed into its corresponding representation. Transformation of input patterns is the process of finding the representation ***a*** that satisfies the equation: ***x*** = ***Φa***. In our case, the dimension of the representational layer is chosen to be higher than that of the input layer, i.e., *K* > *M*. Here, decoding becomes an under-determined problem. Theories developed independently by Donoho (63, 65, 66), and by Candes (44–46) and Tao show that a unique solution can be obtained by imposing a sparseness constraint to the equation when solving the optimization problem. The most common use of sparsity definition includes *L_0_* and *l_1_*. In our approach we perform *L_1_* minimization to solve:

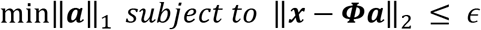

The *l_1_*-minimization problem can be implemented by a standard convex optimization procedure, which can be found in several publications(65, 67–70).

### Redundancy measurement

To measure redundancy in encoding objects, we treated the objects as following a uniform distribution, i.e., 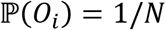, where *N* is the total number of objects. The entropy of the ensemble of the objects is therefore *H*(*O*) = *log N*. We then calculated the capacity of the input unit set (*C*) using the probabilities of occurrence of each encoder, 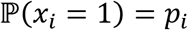:

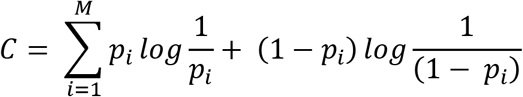

Redundancy was calculated as

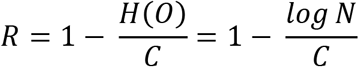

The redundancy for representational units was calculated in a similar way, the only difference being that the representations were converted to binary forms using a Heaviside step function so that their *l_0_* norms could be considered while calculating probability of occurrence of individual encoders.

### KL Divergence between dictionary and images

We used the Kullback–Leibler divergence (KL Divergence) to quantify the structural differences between symbols and dictionary elements. KL divergence 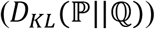 is a measure of information gained when a posterior probability distribution 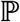 is used to calculate the entropy instead of the prior distribution 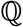. Denoting 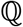 to be distribution over the states of a single pixel in symbol space and 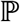 to be the distribution over states of the same pixel in dictionary space, 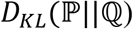 measures the information gained in considering the pixel to be coming from a dictionary element rather than symbols. A low divergence for all the pixels indicates that there is no gain in information if we consider any pixel to be coming from dictionary, indicating that the structure of the dictionary elements is same as structure of the symbols.

To calculate the distribution over the states of pixels in the dictionary space, all dictionary elements were binarized using a Heaviside step function. Probability of occurrence of individual pixels was calculated based on the number of dictionary elements in which the pixel is active. For instance, if a particular pixel *x_i_* was active in *n* out of ***K*** dictionary elements, then the probability of occurrence of pixel *x_i_* was calculated as

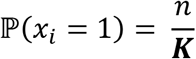

Probability of occurrence of the same pixel in symbol space is calculated based on the number of symbols *m* in which it is active i.e.

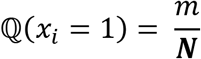

Here *N* is the number of symbols being encoded. Finally, the KL Divergence between the two distributions is calculated as

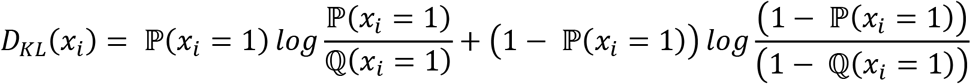

### Specificity calculations

To quantify the specificity of a representational vector in representing the original object, we computed Z-scored similarity. Cosine similarity score between the representation of the test object (***a_test_***) and all objects in the training set (***A_training_***) were calculated and Z-scored. A high Z-score indicated high similarity between the representations of the test object and a particular object in ***A_training_***. In the figures, we plot the Z-scores for altered images with their unadulterated counterpart, which show high specificity in representing the original object.

### Simulating corrupted signals

To test the robustness of object representation by the MDC framework, signals from the training set were selected and corrupted. The corrupted signals were subject to sparse decoding to generate their representations, which were then compared with those of the signals in the training set. We performed the following three types of corruption:

#### Noise-added corruption

We introduce noise by adding a Gaussian i.i.d. matrix 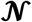 of varying standard deviation to the input matrix ***X***. i.e., 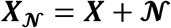, where 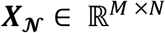, is a matrix representation of noisy input. For Monte Carlo analysis, as described below, each time a simulation was performed, a different noise matrix 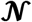 was introduced.

#### Pixel corruption

For a given signals, a fraction of the *M* pixels (glomeruli) was selected from the input. Their values are maintained whereas the coefficients of the rest were set to zero.

#### Occlusion

For images, a contiguous set of pixels were selected, and their coefficient values were set to zero.

### Monte Carlo analysis

We performed Monte Carlo simulations by applying pixel corruption to the input signals and varying the number of corrupted pixels. 100 random sets (numbers varied from 2 to *M*) of pixels (glomeruli) were selected. Using each of these randomly chosen sets, we performed sparse decoding to generate representation of the input patterns.

### Input Identification

To calculate the correct identification of the object, we used the representation of each input in the training set as a library. The representation of each corrupted signal was compared with that in the library and cosine errors were computed. An input pattern was considered correctly identified if the cosine error between its representation and that of the original signal was minimum (smaller than with representation of other patterns).

### Data

#### Symbols

A set of 1000 symbols from word languages were obtained and digitized to 16×16 pixel arrays.

#### Natural images

Natural scenes from Van Hateren data base (49) were digitized as grayscale pictures. Image patches of 16×16 size were randomly selected from the images. A total of 3000 patches were used to form a training set.

#### Facial images

For face recognition, 2000 frontal faces were obtained from Google search of publicly available images, trimmed and resized to 25 x 25 pixels. The Yale Face Database is obtained from http://cvc.cs.yale.edu/cvc/projects/yalefaces/yalefaces.html.

### Projective fields generation from natural images

The mammalian visual systems process visual information in On and Off channels. On channel images were the normal images whereas Off channel images were the inverted images. To simulate parallel processing of the two channels, the On and Off images were concatenated along the rows and dictionary elements were generated by performing BSS on the concatenated matrix. The projective fields were constructed by superposing the On-channel portion of the dictionary element with the negative of Off-channel portion of the dictionary element.

## Supplementary Materials

**Figure S1.**
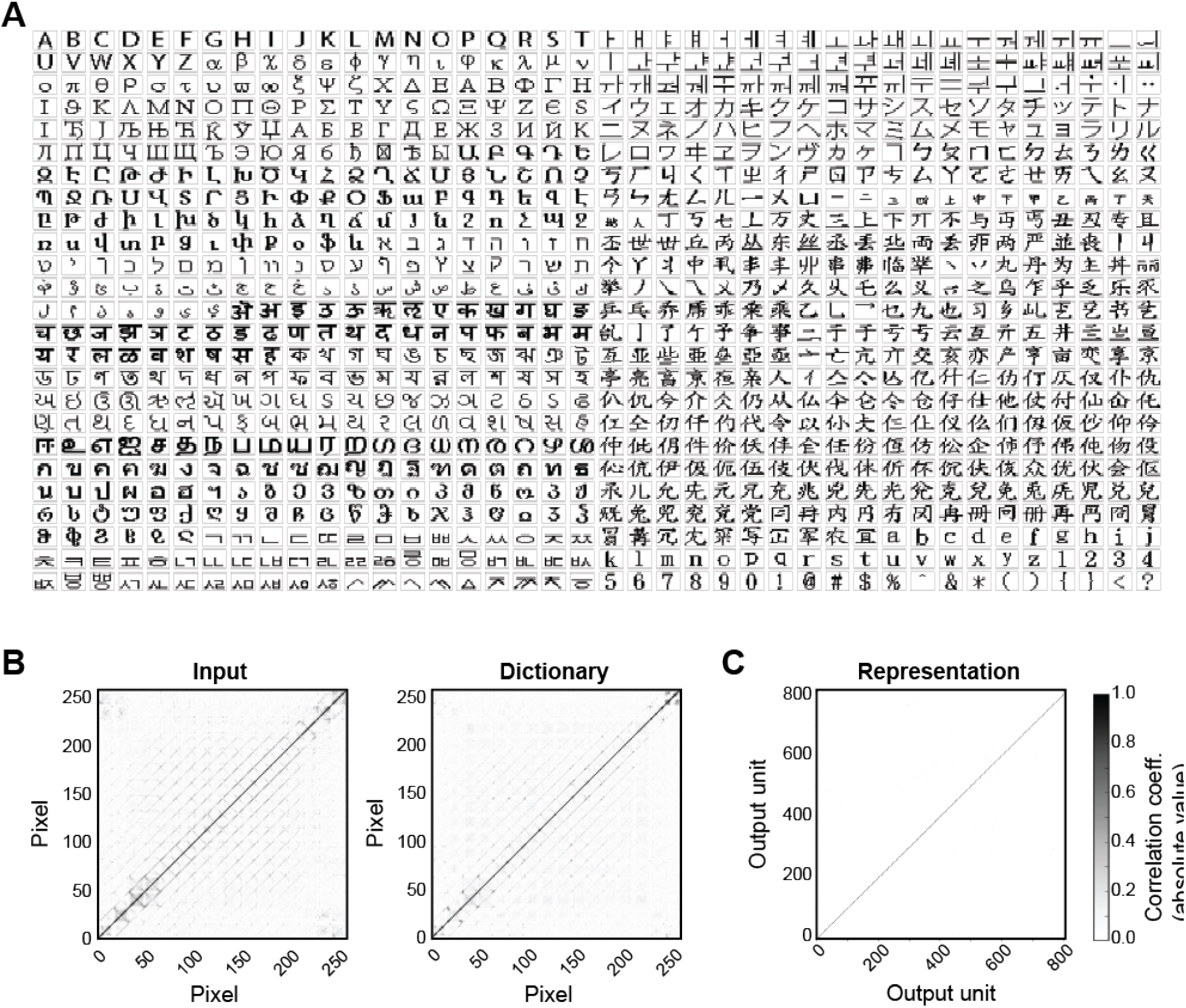
Symbol representations using MDC framework. **(A),** 1000 symbols used in the simulation. **(B),** Correlation between pixels as they occur in symbols and dictionary spaces. **(C),** Correlation between encoding units.

**Figure S2.**
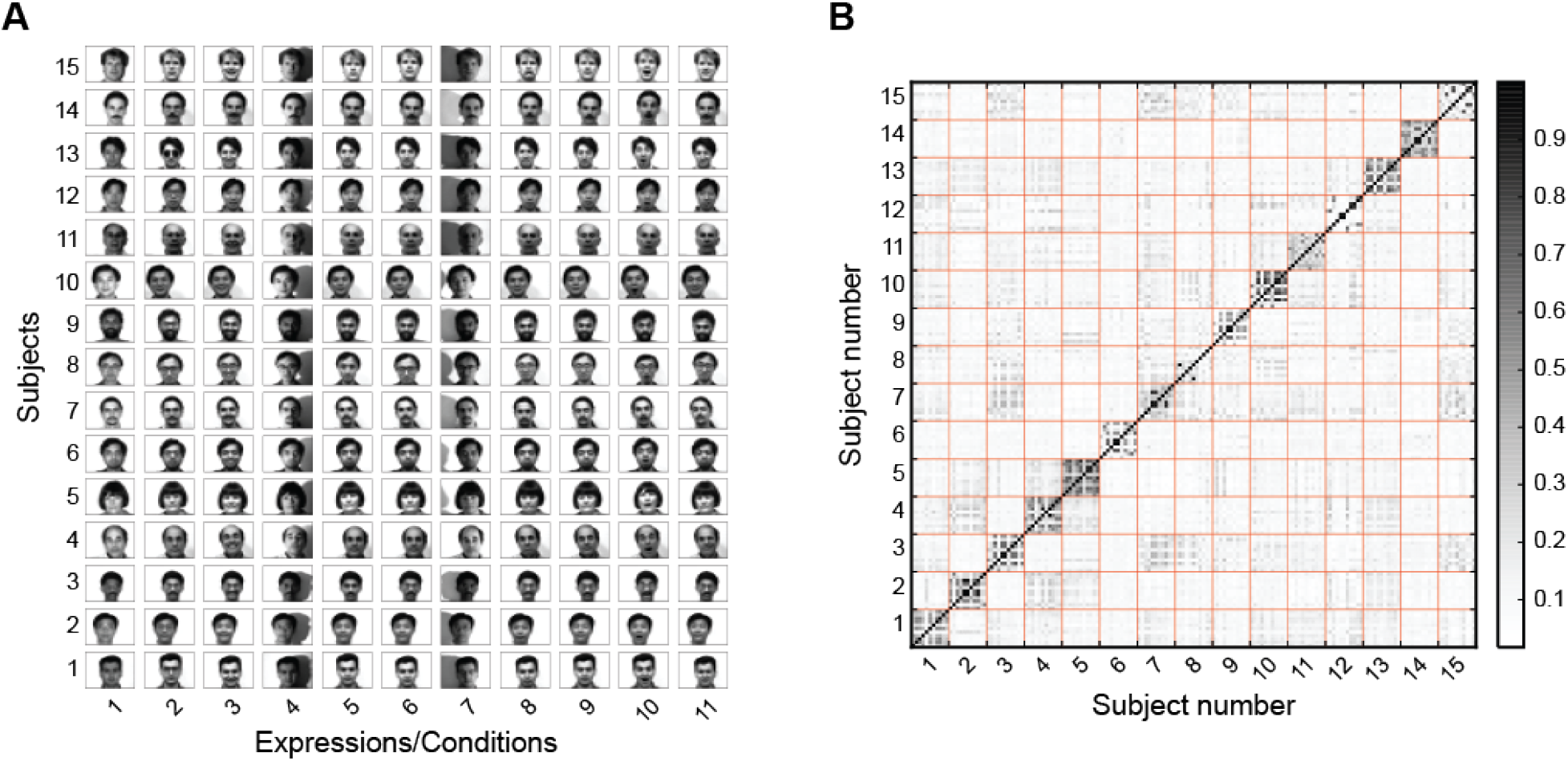
Representation of faces from Yale database. **(A),** images of 15 individuals with different lighting and facial expression. **(B),** Correlation between the output activities sorted according to individuals.

**Figure S3.**
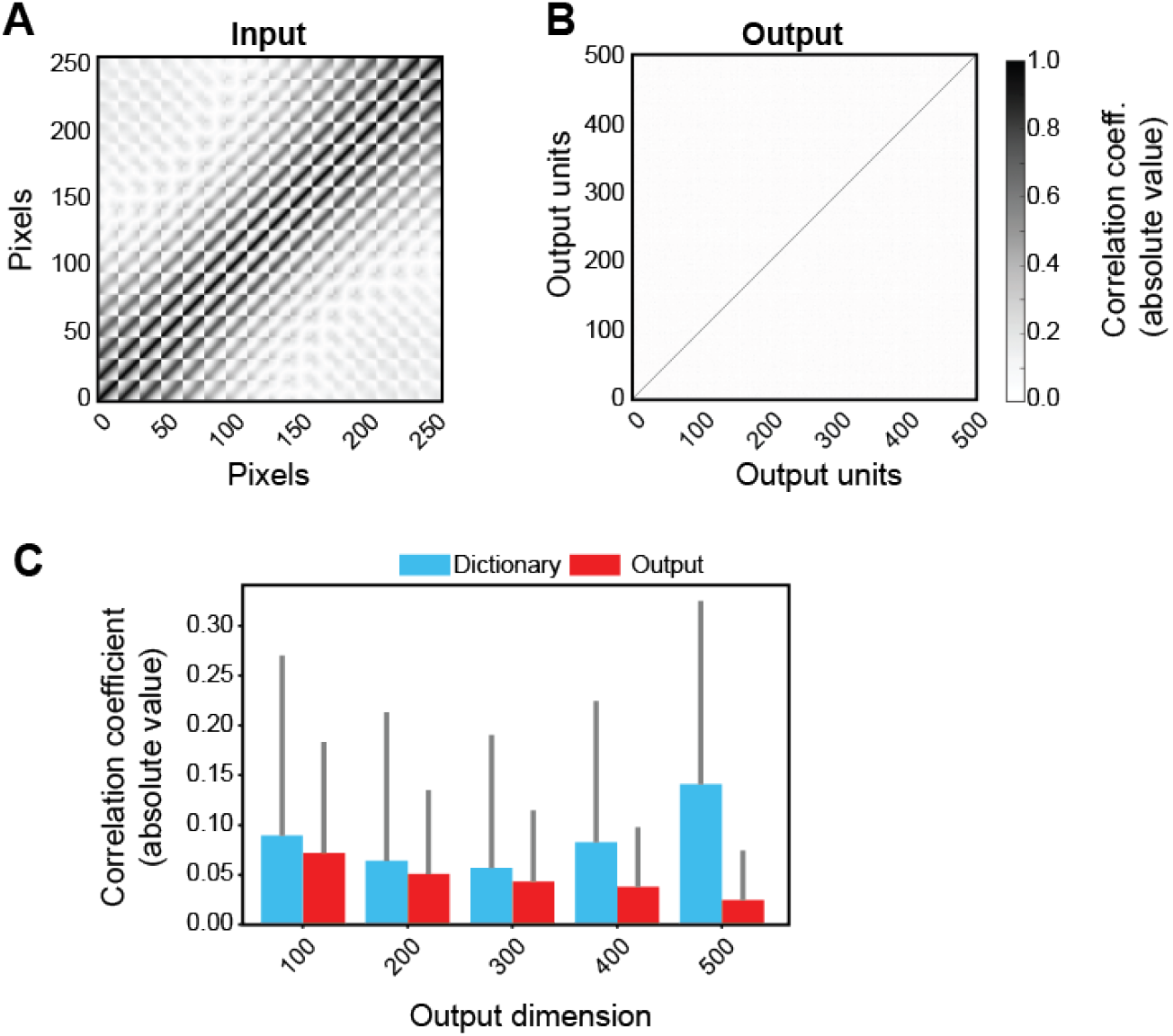
Natural image analysis. **(A),** Correlation among image pixels as they appear in images indicate high levels of redundancy. **(B),** Correlation among output units as they appear in the patches indicate decorrelation among representational units compared to image pixels. **(C),** Correlation among output units decrease as their numbers is increased. Bars indicate the mean of observed correlation coefficients (absolute values) and the whiskers indicate the observed standard deviation in the values.

